## Supplemental Figures for "Dynamic co-evolution of transposable elements and the piRNA pathway in African cichlid fishes"

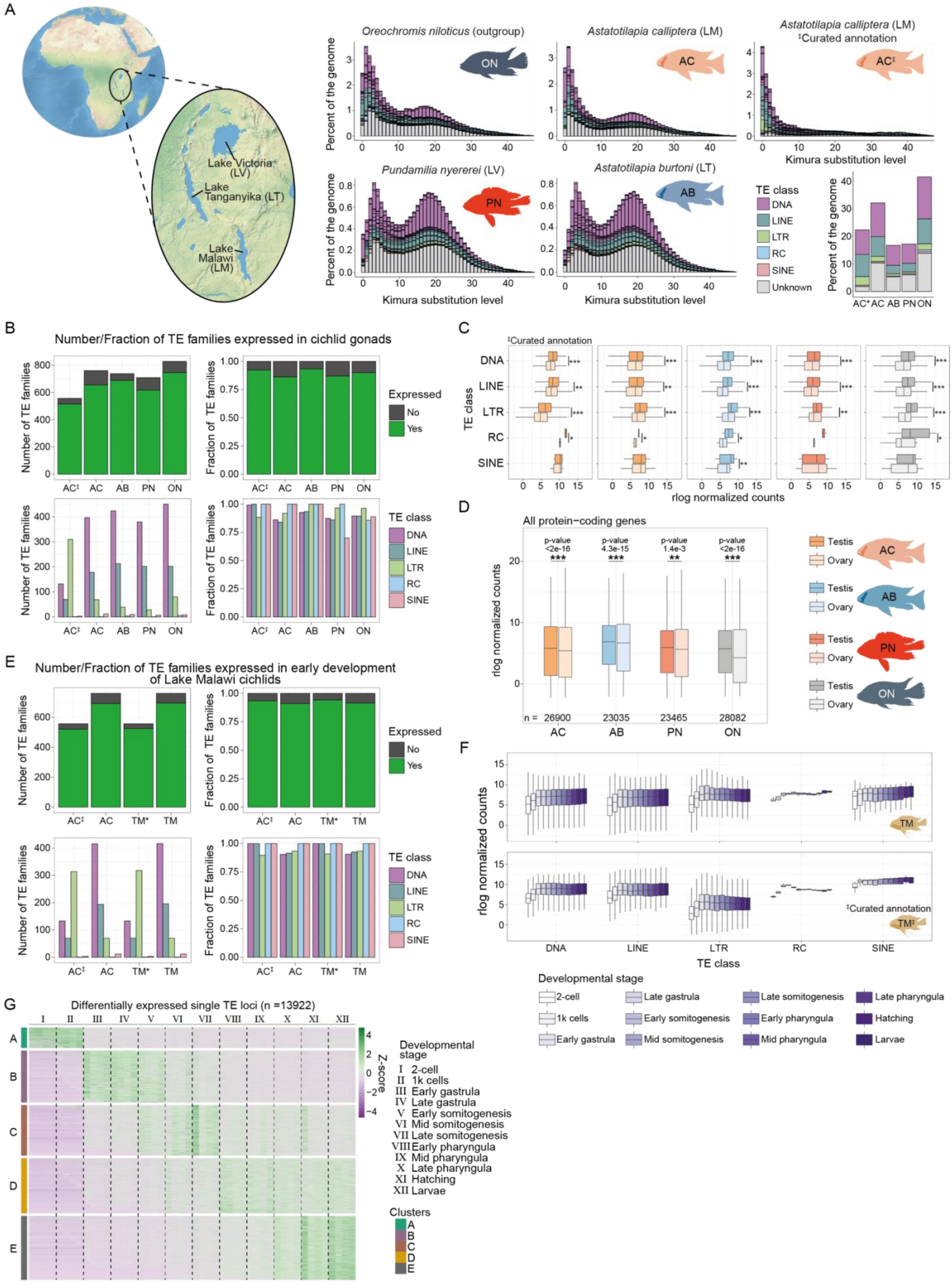

A
Lake Victoria (LV)
Lake Tanganyika (LT)
Lake Malawi (LM)
Oreochromis niloticus (outgroup)
Astatotilapia calliptera (LM)
Astatotilapia calliptera (LM) ‡Curated annotation
Pundamilia nyererei (LV)
Astatotilapia burtoni (LT)
Percent of the genome
Kimura substitution level
TE class
DNA
LINE
LTR
RC
SINE
Unknown
B
Number/Fraction of TE families expressed in cichlid gonads
Number of TE families
Fraction of TE families
Expressed
No
Yes
C
‡Curated annotation
rlog normalized counts
D
All protein−coding genes
p-value <2e-16
p-value 4.3e-15
p-value 1.4e-3
p-value <2e-16
n = 26900 23035 23465 28082
Testis
Ovary
E
Number/Fraction of TE families expressed in early development of Lake Malawi cichlids
F
‡Curated annotation
Developmental stage
2-cell
1k cells
Early gastrula
Late gastrula
Early somitogenesis
Mid somitogenesis
Late somitogenesis
Early pharyngula
Mid pharyngula
Late pharyngula
Hatching
Larvae
G
Differentially expressed single TE loci (n =13922)
Z-score
Developmental stage
I 2-cell
II 1k cells
III Early gastrula
IV Late gastrula
V Early somitogenesis
VI Mid somitogenesis
VII Late somitogenesis
VIII Early pharyngula
IX Mid pharyngula
X Late pharyngula
XI Hatching
XII Larvae
Clusters


**Figure S1.**

**Figure S1. Expression dynamics of TE families and individual TE loci in cichlids.** (**A**) Illustration of the East African Great Lakes as shown in **Figure 1A** and plots depicting the TE landscape of the representative species used in this study. The CpG-adjusted Kimura substitution level is the divergence from a consensus TE modelled for each family. All major TE classes are shown, as well as repeats classified as unknown. For *Astatotilapia calliptera*, the Lake Malawi representative, we use two TE annotations, one of which was curated. The second burst of TE expansion is more prominent in the *P. nyererei* and *A. burtoni* genomes, which have the most fragmented genomes. *O. niloticus* and *A. calliptera* have chromosome-level genome assemblies that display a relatively reduced second burst of TE expansion. Thus, it is likely that genome fragmentation affected TE modelling and TE annotation. A stringently curated TE library (see **Methods**) improved the annotation of younger TE families. (**B**) Number of TE families expressed in cichlid gonads. First two panels depict overall numbers or proportion of expressed families, while the third and fourth panels correspond to the number and proportion of expressed TE families by TE class. (**C**) Expression of TE families in cichlid gonads by TE class, shown as regularised log (rlog) normalised counts. P-values were calculated with Wilcoxon rank-sum tests (using Benjamini & Hochberg correction) comparing expression in ovaries and testes for each TE class. (**D**) Overall expression levels, in rlog normalised counts, of all protein-coding genes in each species in testes versus ovaries. The number of annotated protein-coding genes used for this analysis is indicated above the x axis. P-values were calculated with Wilcoxon rank-sum tests (using Benjamini & Hochberg correction) comparing overall expression in ovaries and testes. (**E**) Same as (B) but for TE families expressed in early development of Lake Malawi cichlids. (**F**) Expression of TE families belonging to major TE classes throughout early development of *Tropheops* sp. 'mauve'. Reads were mapped to the Lake Malawi reference genome (*A. calliptera*) and to non-curated (upper panel) and curated TE annotations (lower panel). (**G**) Heatmap showing differential expression and k-means clustering of individual TE loci in early stages of cichlid development. Analysis done as in Chang et al., 2022[13], using the curated TE annotation of AC. Expression data represented as a z-score. AB, *Astatotilapia burtoni*; AC, *Astatotilapia calliptera*; ON, *Oreochromis niloticus*; PN, *Pundamilia nyererei*; TM, *Tropheops* sp. 'mauve'.

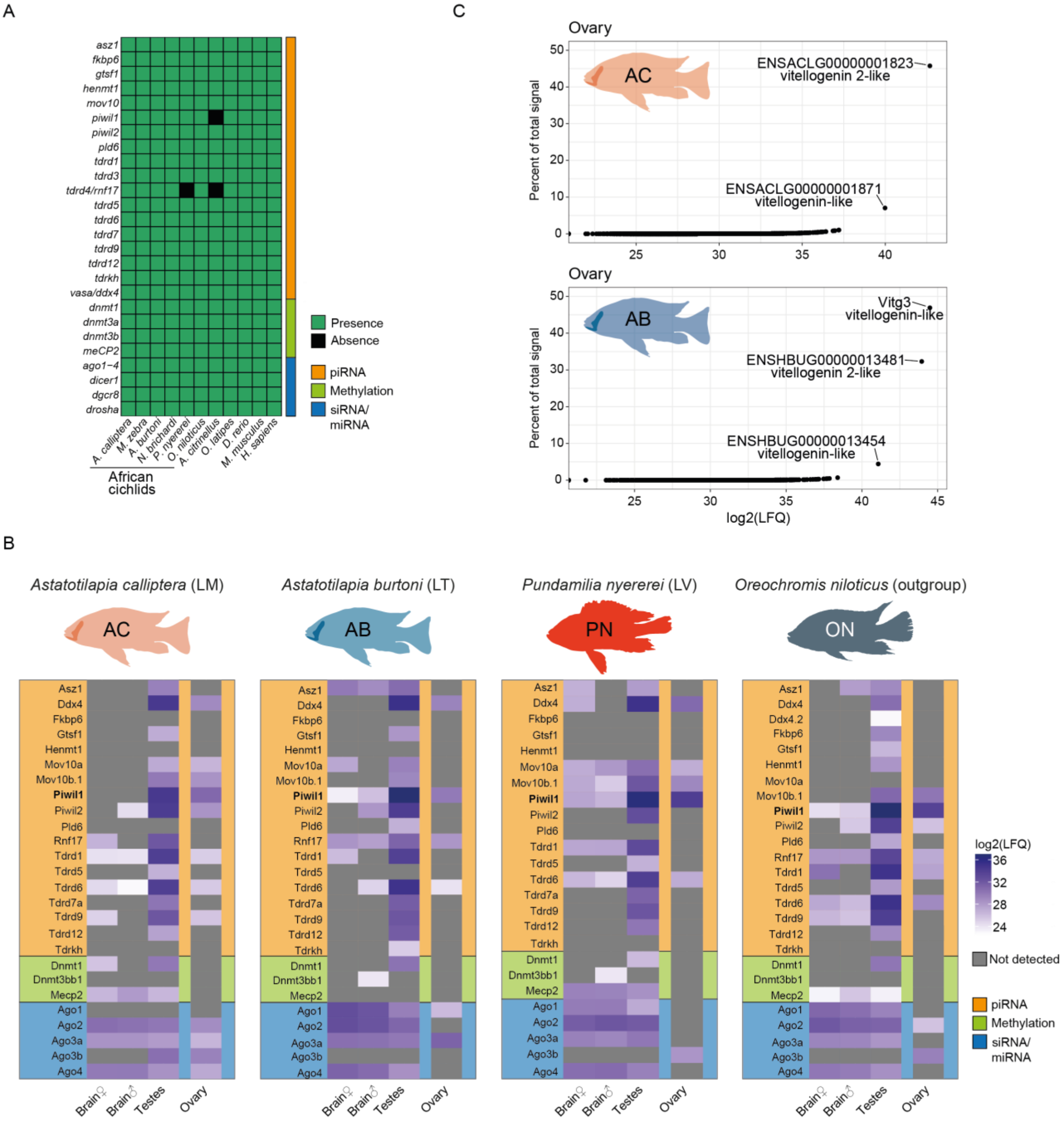

A
C
B
asz1
fkbp6
gtsf1
henmt1
mov10
piwil1
piwil2
pld6
tdrd1
tdrd3
tdrd4/rnf17
tdrd5
tdrd6
tdrd7
tdrd9
tdrd12
tdrkh
vasa/ddx4
dnmt1
dnmt3a
dnmt3b
meCP2
ago1−4
dicer1
dgcr8
drosha
A. calliptera
M. zebra
A. burtoni
N. brichardi
P. nyererei
O. niloticus
A. citrinellus
O. latipes
D. rerio
M. musculus
H. sapiens
African cichlids
Presence
Absence
piRNA
Methylation
siRNA/ miRNA
Ovary
AC
ENSACLG00000001823 vitellogenin 2-like
ENSACLG00000001871 vitellogenin-like
Percent of total signal
AB
Vitg3 vitellogenin-like
ENSHBUG00000013481 vitellogenin 2-like
ENSHBUG00000013454 vitellogenin-like
log2(LFQ)
Astatotilapia calliptera (LM)
Astatotilapia burtoni (LT)
Pundamilia nyererei (LV)
Oreochromis niloticus (outgroup)
PN
ON
Asz1
Ddx4
Ddx4.2
Fkbp6
Gtsf1
Henmt1
Mov10a
Mov10b.1
Piwil1
Piwil2
Pld6
Rnf17
Tdrd1
Tdrd5
Tdrd6
Tdrd7a
Tdrd9
Tdrd12
Tdrkh
Dnmt1
Dnmt3bb1
Mecp2
Ago1
Ago2
Ago3a
Ago3b
Ago4
Brain♀
Brain♂
Testes
Ovary
log2(LFQ)
Not detected


**Figure S2.**

**Figure S2. Conservation and expression of epigenetic silencing factors in African cichlids**. (**A**) Tileplot depicting conservation of epigenetic silencing factors in cichlids. All factors are conserved, besides three exceptions. (**B**) Label-free quantitative proteomics results showing expression of epigenetic silencing factors at the protein level. Only proteins detected at least in one organ in one species are shown. Proteins not detected in any of the species (TDRD3, TDRD7B, DNMT3AA, DNTM3AB, DNMT3BB2, DICER1, DGCR8, and DROSHA) are not shown. Ovary samples are shown in an isolated column to depict that direct comparisons with the other organs are to be avoided, as protein detection in ovaries was hampered by extremely abundant yolk proteins. (**C**) Extremely abundant yolk proteins are detected in label-free quantitative proteomics of ovaries (shown as percent of total LFQ signal), precluding robust detection of other proteins. AB, *Astatotilapia burtoni*; AC, *Astatotilapia calliptera*; LM, Lake Malawi; LT, Lake Tanganyika; LV, Lake Victoria; ON, *Oreochromis niloticus*; PN, *Pundamilia nyererei*.

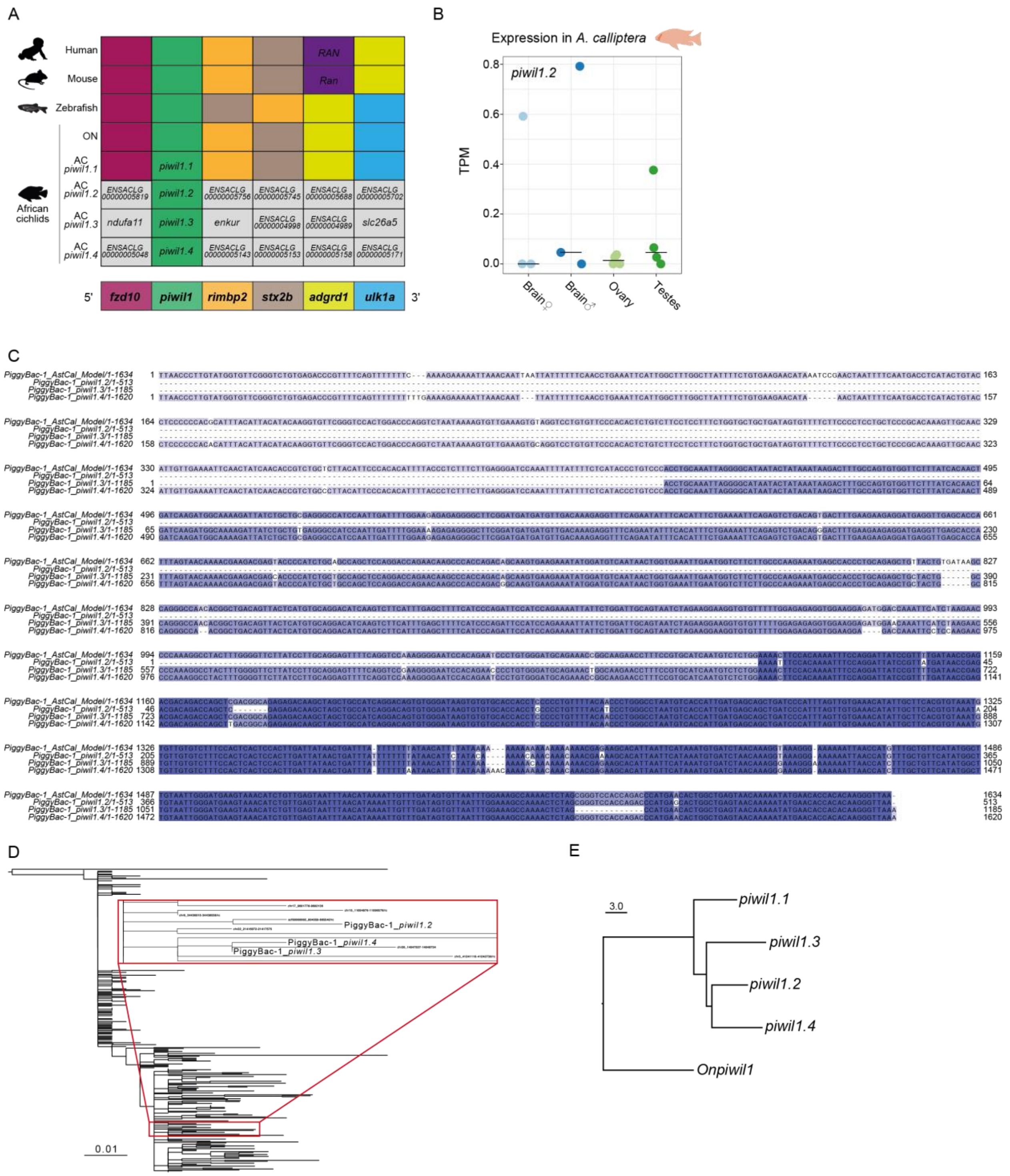

A
Human
Mouse
Zebrafish
ON
AC piwil1.1
AC piwil1.2
AC piwil1.3
AC piwil1.4
African cichlids
RAN
Ran
piwil1.1
piwil1.2
piwil1.3
piwil1.4
ndufa11
enkur
slc26a5
5'
fzd10
piwil1
rimbp2
stx2b
adgrd1
ulk1a
3'
B
Expression in A. calliptera
piwil1.2
TPM
Brain♀
Brain♂
Ovary
Testes
C
PiggyBac-1_AstCal_Model/1-1634
PiggyBac-1_piwil1.2/1-513
PiggyBac-1_piwil1.3/1-1185
PiggyBac-1_piwil1.4/1-1620
D
PiggyBac-1_piwil1.2
PiggyBac-1_piwil1.4
PiggyBac-1_piwil1.3
0.01
E
3.0
piwil1.1
piwil1.3
piwil1.2
piwil1.4
Onpiwil1


**Figure S3.**

**Figure S3. Additional data on the expansion of *piwil1* genes in Lake Malawi cichlids**. (**A**) Synteny analysis of *piwil1* genes in vertebrates. Each row of adjacent squares represents the order, oriented from 5' to 3', of genes adjacent to *piwil1* (in green). Squares of the same colour represent the same gene. The lower row of squares shows the consensus synteny and the correspondence between the colour-code and each gene. Grey squares indicate genes in linkage with only one *piwil1* gene. Of all *A. calliptera piwil1* genes, *piwil1.1* is the only gene syntenic with the *piwil1* gene of other vertebrates. AC, *Astatotilapia calliptera*; ON, *Oreochromis niloticus*. (**B**) Expression of *piwil1.2* in gonads and brain of *A. calliptera*, in Transcripts per Million (TPM). (**C**) Multiple sequence alignment of the PiggyBac-1 family consensus model of the curated *A. calliptera* TE library (sequence at the top) with the PiggyBac-1 sequences directly 3' of *piwil1.2*, *piwil1.3*, and *piwil1.4*. Colouring is according to sequence identity. According to open reading frame predictions, the PiggyBac-1 family is not expected to encode a full-length transposase. Additional mutations have accumulated in *piwil1*-associated PiggyBac-1 TEs. (**D**) Phylogenetic tree constructed from all PiggyBac-1 TEs in the *A. calliptera* genome that align with piwil1-associated PiggyBac-1 TEs. Tree was rooted at the midpoint. The branches with the *piwil1*-associated PiggyBac-1 TEs are shown in detail in the red box inset, illustrating their relatedness. (**E**) Neighbour-joining tree representing the Hamming distance between the coding sequences within the region shared by all *A. calliptera piwil1* genomic sequences (Region S in **Figure 2A**). The equivalent region in *piwil1* of *O. niloticus* (*Onpiwil1*) is included as an outgroup.

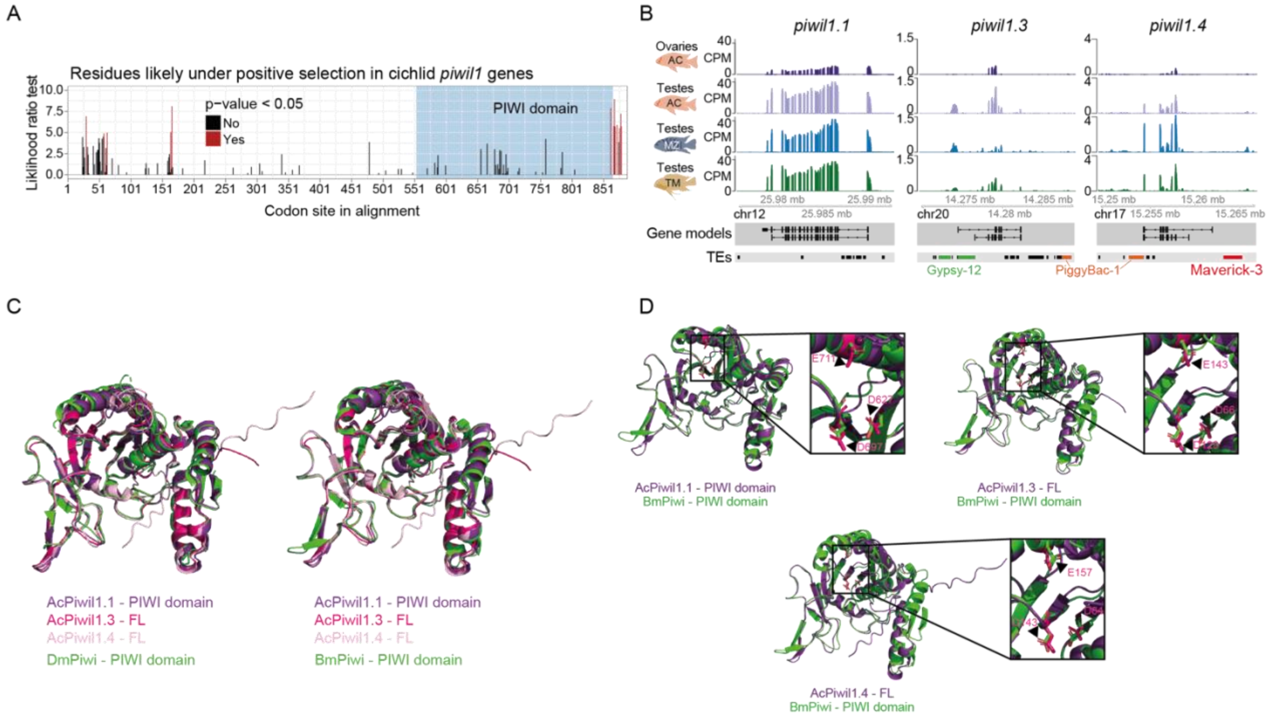
A
Residues likely under positive selection in cichlid piwil1 genes
Liklihood ratio test
p-value < 0.05
No
Yes
PIWI domain
Codon site in alignment
B
piwil1.1
piwil1.3
piwil1.4
Ovaries
Testes
CPM
Gene models
TEs
Gypsy-12
PiggyBac-1
Maverick-3
C
AcPiwil1.1 - PIWI domain
AcPiwil1.3 - FL
AcPiwil1.4 - FL
DmPiwi - PIWI domain
AcPiwil1.1 - PIWI domain
AcPiwil1.3 - FL
AcPiwil1.4 - FL
BmPiwi - PIWI domain
D
AcPiwil1.1 - PIWI domain
BmPiwi - PIWI domain
AcPiwil1.3 - FL
BmPiwi - PIWI domain
AcPiwil1.4 - FL
BmPiwi - PIWI domain

**Figure S4.**

**Figure S4. Expression, function, and evolution of cichlid *piwil1* genes**. (**A**) Residues very likely to be under positive selection according to Mixed Effects Model of Evolution (MEME)[92] across the entire alignment of African cichlid Piwil1 coding sequences. A region in the C-terminal portion of the proteins has 8 amino acid residues predicted to be under positive selection. (**B**) Genome tracks showing the mRNA expression of *piwil1.1* (left panel), *piwil1.3* (central panel), and *piwil1.4* (right panel). Large TE fragments flanking *piwil1.3* and *piwil1.4* are annotated and coloured in the TE track. mRNA expression shown in Counts per Million (CPM). (**C**) Structural alignments of the AlphaFold-predicted PIWI domain of *Astatotilapia calliptera* (Ac) Piwil1.1, or full-length protein in case of Piwil1.3 and Piwil1.4, with the PIWI domain of published crystal structures of *Drosophila melanogaster* (Dm) Piwi (left) and *Bombyx mori* (Bm) Siwi (right). (**D**) Structural alignments of the PIWI domain of Bombyx mori (Bm) Siwi protein with the AlphaFold predictions of the Piwil1.1 (using only PIWI domain), Piwil1.3 (full-length), and Piwil1.4 (full-length) of *A. calliptera*. Insets focus on the regions with the integral residues of the catalytic triad, which are indicated with black or white arrowheads. AC, *Astatotilapia calliptera*; MZ, *Maylandia zebra*; TM, *Tropheops* sp. ‘mauve’.

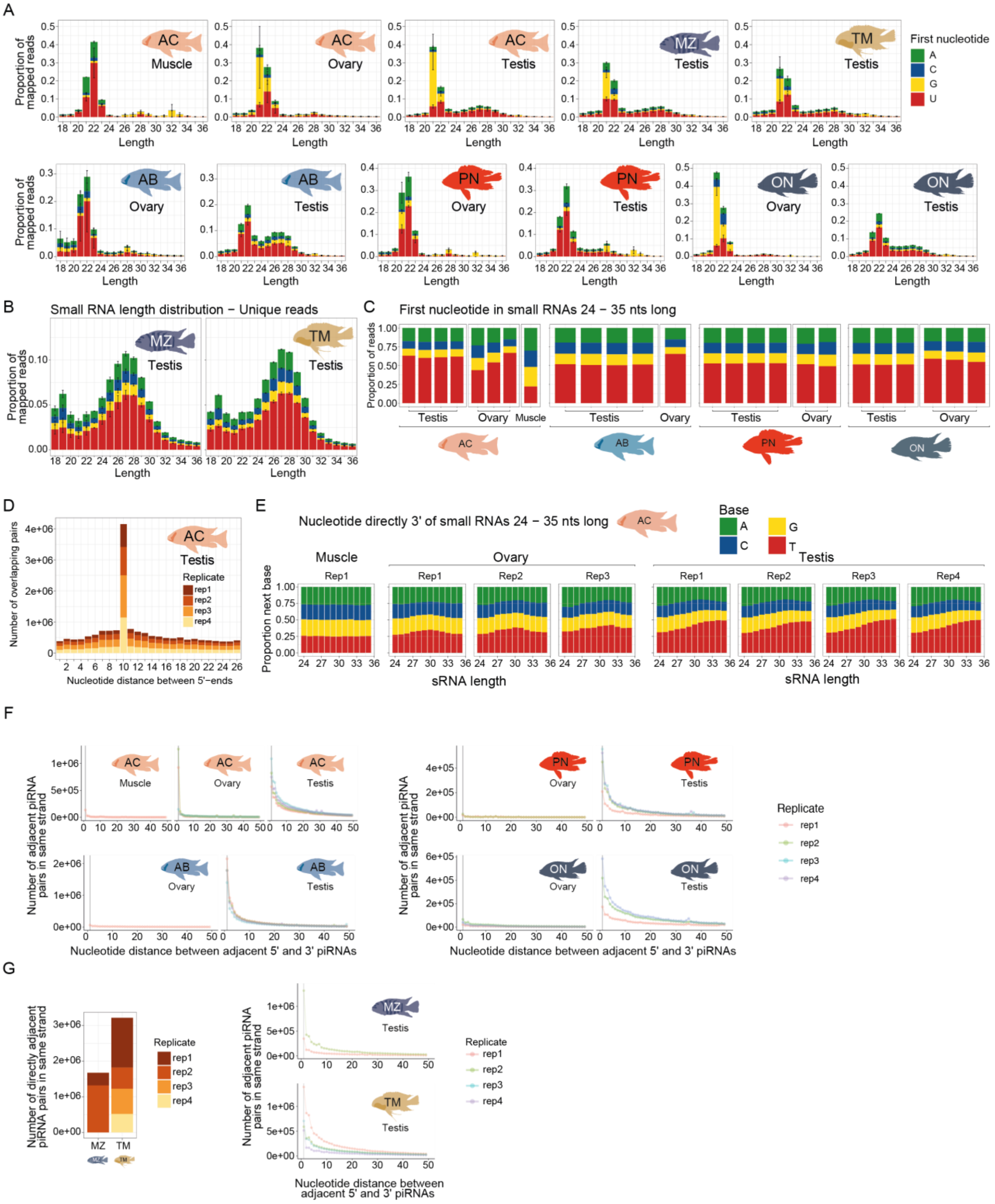
A
Proportion of mapped reads
Length
AC Muscle
AC Ovary
AC Testis
MZ Testis
TM Testis
First nucleotide
A
C
G
U
AB Ovary
AB Testis
PN Ovary
PN Testis
ON Ovary
ON Testis
B
Small RNA length distribution − Unique reads
MZ Testis
TM Testis
C
First nucleotide in small RNAs 24 − 35 nts long
Proportion of reads
Testis
Ovary
Muscle
AC
AB
PN
ON
D
Number of overlapping pairs
AC Testis
Replicate
rep1
rep2
rep3
rep4
Nucleotide distance between 5'−ends
E
Nucleotide directly 3' of small RNAs 24 − 35 nts long
Base
A
C
G
T
Muscle
Ovary
Testis
Rep1
Rep2
Rep3
Rep4
Proportion next base
sRNA length
F
Number of adjacent piRNA pairs in same strand
Nucleotide distance between adjacent 5' and 3' piRNAs
Replicate
rep1
rep2
rep3
rep4
G
Number of directly adjacent piRNA pairs in same strand
MZ
TM
Nucleotide distance between adjacent 5' and 3' piRNAs

**Figure S5.**

**Figure S5. Profile and sequence signatures of sRNAs in African cichlids.** (**A**) sRNA length distribution profiles of all reads, without collapsing into unique reads, display prominent peaks at 21-22 nucleotides, likely attributable to abundant microRNAs. Colours display the identity of the first nucleotide in the sRNA. (**B**) sRNA length distribution profiles of unique reads in *M. zebra* and *T.* sp. ‘mauve’ testes. Identity of first nucleotide is indicated by the colouring, which is colour coded as in (A). (**C**) First nucleotide identity in sRNAs 24-35 nucleotides long. Same colouring by first nucleotide as in (A). (**D**) Number of overlapping sRNA read pairs according to the length of the overlap in *A. calliptera* testis replicates. The peak at 10 nucleotides supports a ping-pong signature. (**E**) Identity of nucleotide directly 3' of the last nucleotide of sRNAs 24-35 nucleotides long. A bias for a T is consistent with a signature of phased piRNA biogenesis. (**F**) Number of directly adjacent piRNA pairs and the distance separating them. Grey line indicates distance of 1 nucleotide, indicative of phased piRNA biogenesis, where two piRNAs are produced consecutively. Testes of all species have high numbers of piRNA pairs at 1 nucleotide distance, but this signature is not clear in ovaries, except the ovaries of AC. (**G**) On the right, plots as in (F) showing the number of piRNA pairs and the distance between each member of the pair. Grey line marks distance of 1 nucleotide. On the left, histogram with the number of piRNA pairs distant 1 nucleotide from each other (equivalent to the grey line in the right-hand side plots). AB, *Astatotilapia burtoni*; AC, *Astatotilapia calliptera*; MZ, *Maylandia zebra*; ON, *Oreochromis niloticus*; PN, *Pundamilia nyererei*; TM, *Tropheops* sp. ‘mauve’; sRNA, small RNA.

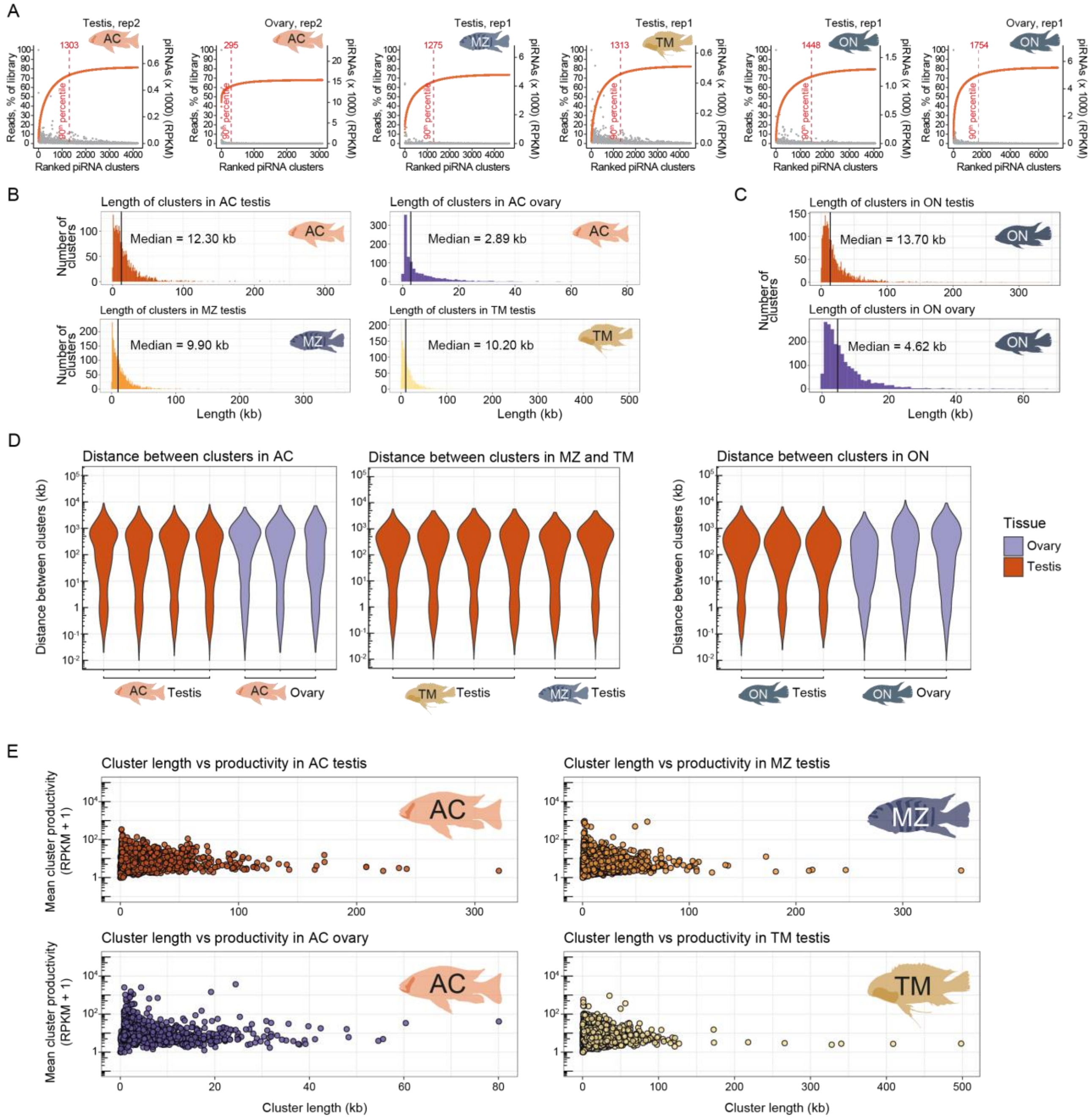
A
Testis, rep2
Ovary, rep2
Testis, rep1
Testis, rep1
Testis, rep1
Ovary, rep1
Reads, % of library
piRNAs (x 1000) (RPKM)
90th percentile
Ranked piRNA clusters
B
Length of clusters in AC testis
Median = 12.30 kb
Length of clusters in AC ovary
Median = 2.89 kb
Length of clusters in MZ testis
Median = 9.90 kb
Length of clusters in TM testis
Median = 10.20 kb
Number of clusters
Length (kb)
C
Length of clusters in ON testis
Median = 13.70 kb
Length of clusters in ON ovary
Median = 4.62 kb
D
Distance between clusters in AC
Distance between clusters in MZ and TM
Distance between clusters in ON
Distance between clusters (kb)
Tissue
Ovary
Testis
E
Cluster length vs productivity in AC testis
Cluster length vs productivity in MZ testis
Cluster length vs productivity in AC ovary
Cluster length vs productivity in TM testis
Mean cluster productivity (RPKM + 1)
Cluster length (kb)

**Figure S6.**

**Figure S6. Additional features of cichlid piRNA clusters**. (**A**) Ranking of piRNA clusters according to productivity across Lake Malawi cichlids and *O. niloticus*. Cumulative fraction shown as an orange line. Dashed red line indicates the 90$^{th}$ percentile of piRNA production and the number of clusters producing 90% of piRNA reads in clusters. (**B**) Histograms showing the length distribution of piRNA clusters in *A. calliptera* (two upper panels) *M. zebra* (lower left panel) and *T.* sp. 'mauve' (lower right panel). Black line indicates the median. (**C**) Histogram showing the distribution of the lengths of piRNA clusters in *O. niloticus*. Black line indicates the median. (**D**) Distance between clusters, in kilobase (kb), in each replicate of the indicated organs and species. Left panel for *A. calliptera* gonads, central panel for *T.* sp. 'mauve' and *M. zebra* testes, and right panel for *O. niloticus* gonads. (**E**) Relationship between piRNA cluster length and cluster productivity. Productivity is plotted as mean Reads Per Kilobase Million (RPKM) of a cluster across all replicates. AC, *Astatotilapia calliptera*; MZ, *Maylandia zebra*; ON, *Oreochromis niloticus*; TM, *Tropheops* sp. 'mauve'.

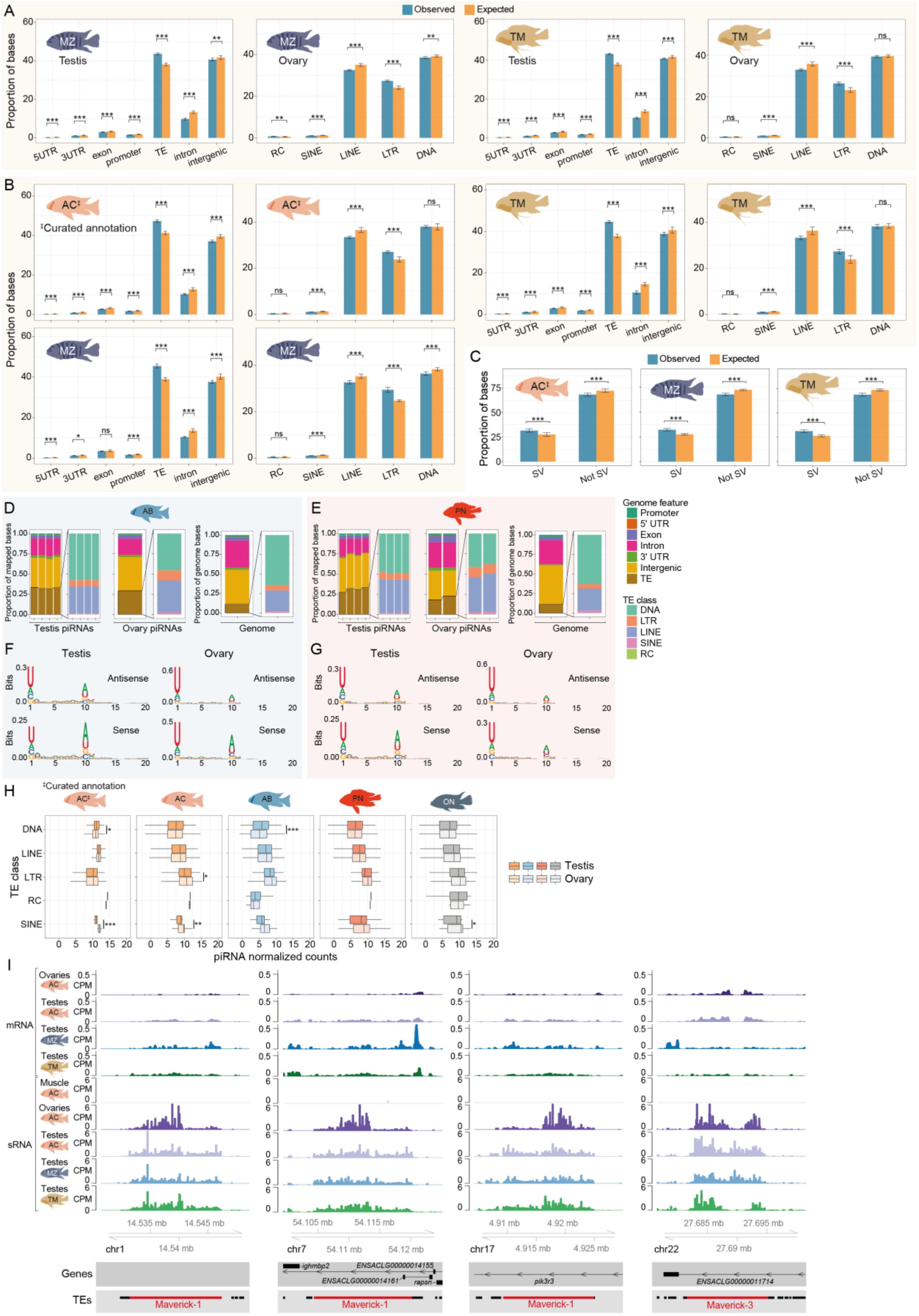
A
Observed
Expected
Proportion of bases
MZ
Testis
Ovary
TM
5UTR
3UTR
exon
promoter
TE
intron
intergenic
RC
SINE
LINE
LTR
DNA
B
AC‡
‡Curated annotation
C
SV
Not SV
D
AB
Proportion of mapped bases
Proportion of genome bases
Testis piRNAs
Ovary piRNAs
Genome
E
PN
Genome feature
Promoter
5' UTR
Exon
Intron
3' UTR
Intergenic
TE
TE class
DNA
LTR
LINE
SINE
RC
F
Antisense
Sense
Bits
G
H
AC
ON
TE class
Testis
Ovary
piRNA normalized counts
I
mRNA
sRNA
Ovaries
Testes
Muscle
CPM
14.535 mb
14.545 mb
chr1
14.54 mb
54.105 mb
54.115 mb
chr7
54.11 mb
54.12 mb
4.91 mb
4.92 mb
chr17
4.915 mb
4.925 mb
27.685 mb
27.695 mb
chr22
27.69 mb
Genes
TEs
ighmbp2
ENSACLG00000014155
ENSACLG00000014161
rapsn
pik3r3
ENSACLG00000011714
Maverick-1
Maverick-3

**Figure S7.**

**Figure S7. The genomic origin and sequence signatures of piRNAs targeting TEs in cichlids**. (**A**) Observed and expected values at genomic features and TE classes that piRNA clusters overlap with in Lake Malawi cichlids MZ and TM. (**B**) The genomic composition of the diverging piRNA clusters in Lake Malawi cichlids, present in one species (AC, MZ, or TM), but not in at least one of the other species. The species where the diverging clusters is present is indicated in the panel. In (B), TE features used to calculate overlaps with piRNA clusters correspond to the curated TE annotation. (**C**) Proportion of mapped bases of species-variable piRNA clusters that overlap with structural variants (SVs) in the Lake Malawi pangenome, defined in Quah et al., 2024[5]. The species where the variable clusters are present is indicated in the panel. (**D-E**) Genomic features that 24-35 nucleotide long piRNAs map to in *A. burtoni* (D) and *P. nyererei* (E). Each bar represents a separate replicate. Inset barplots specify the proportions of major TE classes that piRNAs map to. Genome bars represent the proportion of bases corresponding to genome-wide specific features and TE classes. (**F-G**) Sequence logos of 24-35 nucleotide long piRNAs mapping sense or antisense in regard to TE orientation in *A. burtoni* (F) and *P. nyererei* (G). (**H**) Expression of piRNAs mapping to the major TE classes. P-values were calculated with Wilcoxon rank-sum tests (using Benjamini & Hochberg correction) comparing expression in ovaries and testes for each TE class. We note that incomplete LTR annotation may overestimate the expression of LTR-mapping piRNAs (compare AC curated versus non-curated annotation). (**I**) Genome tracks with the mRNA expression and 24-35 nucleotide sRNAs mapping to large Maverick TEs in Lake Malawi cichlids. mRNA and sRNA expression shown in Counts per Million (CPM). In the TE track, a curated TE annotation is shown, with the large Maverick element coloured in red. AB, *Astatotilapia burtoni*; AC, *Astatotilapia calliptera*; CPM, Counts per Million; MZ, *Maylandia zebra*; ns, not statistically significant; PN, *Pundamilia nyererei*; SV, structural variant; TM, *Tropheops* sp. 'mauve'; sRNA, small RNA.
